# Intranasal Nanoemulsion Adjuvanted S-2P Vaccine Demonstrates Protection in Hamsters and Induces Systemic, Cell-Mediated and Mucosal Immunity in Mice

**DOI:** 10.1101/2022.03.22.485323

**Authors:** Shyamala Ganesan, Hugo Acosta, Chris Brigolin, Kallista Orange, Kevin Trabbic, Charles Chen, Chia-En Lien, Yi-Jiun Lin, Meei-Yun Lin, Ya-Shan Chuang, Ali Fattom, Vira Bitko

## Abstract

With the rapid progress made in the development of vaccines to fight the SARS-CoV-2 pandemic, almost >90% of vaccine candidates under development and a 100% of the licensed vaccines are delivered intramuscularly (IM). While these vaccines are highly efficacious against COVID-19 disease, their efficacy against SARS-CoV-2 infection of upper respiratory tract and transmission is at best temporary. Development of safe and efficacious vaccines that are able to induce robust mucosal and systemic immune responses are needed to control new variants. In this study, we have used our nanoemulsion adjuvant (NE01) to intranasally (IN) deliver stabilized spike protein (S-2P) to induce immunogenicity in mouse and hamster models. Data presented demonstrate the induction of robust immunity in mice resulting in 100% seroconversion and protection against SARS-CoV-2 in a hamster challenge model. There was a significant induction of mucosal immune responses as demonstrated by IgA- and IgG-producing memory B cells in the lungs of animals that received intranasal immunizations compared to an alum adjuvanted intramuscular vaccine. The efficacy of the S-2P/NE01 vaccine was also demonstrated in an intranasal hamster challenge model with SARS-CoV-2 and conferred significant protection against weight loss, lung pathology, and viral clearance from both upper and lower respiratory tract. Our findings demonstrate that intranasal NE01-adjuvanted vaccine promotes protective immunity against SARS-CoV-2 infection and disease through activation of three arms of immune system: humoral, cellular, and mucosal, suggesting that an intranasal SARS-CoV-2 vaccine may play a role in addressing a unique public health problem and unmet medical need.

## Introduction

The respiratory virus causing COVID-19 is a zoonotic betacoronavirus known as SARS-CoV-2 (Severe Acute Respiratory Syndrome Coronavirus 2).^1^ SARS-CoV-2 is an enveloped positive-sense RNA virus related to the previous coronavirus infections caused by Middle East Respiratory Syndrome MERS-CoV and SARS-CoV with ~50% and ~80 % nucleotide sequence identity, respectively.

SARS-CoV-2 infection is predominantly initiated by entry of aerosolized respiratory droplets to the upper respiratory tract (URT) through the nasal passages.^2^ In the nasal passages, the viral spike protein (S) facilitates entry into cells by binding to the angiotensin-converting enzyme 2 (ACE2) receptor. The S protein is a trimeric glycoprotein (180-200 kDa) whose ectodomain is composed of two subunits, S1 and S2. The S1 subunit contains the receptor-binding domain (RBD). The S2 subunit is responsible for initiating the viral-host membrane fusion and is activated by cleavage of the pre-fusion protein through transmembrane protease serine 2 (TMPRS2). Increased viral load and subsequent host dissemination is supported by elevated levels of ACE2 co-expressed with TMPRS2 in nasal ciliated cells and localization in the URT.^3–5^ Viral seeding in the nasal cavity supports efficient initiation of infection and propagation of the virus to a high viral titer prior to inoculation of the lungs and initiation of SARS-CoV-2 infection cascade.^5–7^ The URT infection phase is the most infectious and, key in disseminating viral spread. Developing a vaccine to induce mucosal immunity at the port of viral entry will prevent viral colonization and prevent subsequent infection of the lungs and disease transmission.

The introduction of highly effective SARS-CoV-2 vaccines early in the pandemic has curbed the pandemic and saved millions of human lives.^8, 9^ Different technologies, new and old, were employed to develop these vaccines. The first approach utilizes mRNA delivery systems produced by Pfizer-BioNTech (BNT162b2) and Moderna (mRNA-1273). Both mRNA vaccines consist of lipid nanoparticle encapsulating modified mRNA encoding a stable prefusion S protein. The BNT162b2 and mRNA-1273 vaccines reported a vaccine efficacy of ~95% and ~94%, respectively, after the administration of two doses.^10, 11^ The second vaccine approach produced by Johnson & Johnson (Ad26.COV2.S) and Oxford/AstraZeneca ^12, 13^ (AZD1222/ChAdOx1 nCoV-19) use adenoviral vectors encoding the spike protein. ‘ Collectively, all these vaccines are highly effective in inducing protective neutralizing antibodies in serum, thereby preventing severe COVID-19 disease, which leads to hospitalizations. However, these highly efficacious intramuscularly delivered vaccines only provide partial protection against URT infection and transmission of the virus, which possibly added to the subsequent waves of SARS-CoV-2 variant infections.^14, 15^ Only intranasal vaccines capable of inducing mucosal immunity can prevent nasal infection, shedding and further transmission of the virus effectively, via localized URT immunity and memory responses in addition to inducing systemic immune responses.

Intranasal immunization introduces antigens to immune cells that will process and drain them to the nasal-associated lymphoid tissue (NALT). Subsequently, a mucosal and systemic immune response is elicited that includes secretory IgA and IgG, homing of B and T cells to mucosal tissues, and induction Th17 cells. Serum neutralizing and systemic memory B and T-cells are also induced. Induction of both mucosal and systemic immunity are essential for complete protection against infection, disease, and spread to others.

In this study, we employed mouse and hamster models to evaluate an intranasal S-2P nanoemulsion-adjuvanted vaccine to generate a complete and potent immune response to SARS-CoV-2 that protects against colonization, viral spread, and disease.

## Materials and Methods

### SARS-CoV-2S (S-2P)protein

Recombinant stabilized trimeric full length S protein was provided by Medigen Vaccine Biologics Corporation. The SARS-CoV-2 (Wuhan-Hu-1 strain, GenBank: MN908947) S-2P protein contains the residues 1-1208 with a C-terminal T4 fibritin trimerization domain, an HRV3C cleavage site, an 8X His-tag and a Twin-Strep-tag. The stabilized S-2P form was achieved by mutation of the S1/S2 furin-recognition site 682-RRAR-685 to GSAS to produce a single chain S protein, and the 986-kV-987 was mutated to PP. The protein was produced in Expi-CHO-S cells as described previously. ^16, 17^

### Nanoemulsion Adjuvant and Vaccine Preparation

The 60% NE01 was prepared by high shear homogenization of water, ethanol, cetylpyridinium chloride, Tween-80 (non-ionic surfactant), and highly refined soybean oil to form an oil-in-water nanoemulsion with a mean particle size of ~400 nm as described previously.^18^ The vaccine was prepared by mixing S-2P with NE01 adjuvant for a final concentration of 2.5 μg of S-2P (mouse studies) or 10 μg of S-2P (hamster studies) with 20% NE01/dose.

Alum adsorbed intramuscular vaccine was prepared by mixing 2.5 μg of S-2P with 30 μg of alum (Croda, Cat# AJV3012) in a 50 μL dose volume. The prepared vaccine was mixed thoroughly before administering to animals.

### Mouse Study

Mouse immunization studies were performed at IBT Bioservices, Rockville, MD, USA under the approved IACUC animal study protocol # AP-160805. Six-to eight-week-old female CD-1 mice were randomly assigned to each of the five groups, with 8 animals in each, except for group with two intranasal vaccinations, where 7 animals were assigned. Mice were immunized intranasally with S-2P/NE01 either three or two times, or three times with S-2P alone, or intramuscularly with S-2P/alum, and an unimmunized control group. All vaccinated animals received 2.5 μg of S-2P protein/dose either in 12 μL (intranasal dose), or 50 μL (intramuscular dose). Vaccines were administered three weeks apart and blood was collected two weeks post last vaccination. Bronchio-alveolar lavage (BAL) was collected prior to collection of lungs on week 8 (day 56), followed by collection of lungs and spleens.

### Hamster Study

Hamster challenge studies were performed at Testing Facility for Biological Safety, TFBS Bioscience Inc., Taiwan and Academia Sinica, Taiwan. Six-to nine-week-old female golden Syrian hamsters were randomized into four groups. Groups of 12 hamsters were immunized either with three or one intranasal dose, while the group vaccinated two times had 10 animals. There were six animals assigned to the negative control group (PBS vaccination). Hamsters were immunized three weeks apart with 10μg/20μL (10μL/nare) of S-2P/NE01 per each dose. Animals were challenged with SARS-CoV-10 days post last dose (as described below) and finally they were bled three weeks after the last vaccine dose. The studies were performed with approval by the IACUC with animal study protocol approval number TFBS2020-019 and Academia Sinica (approval number:20-10-1526)

### Hamster challenge with SARS-CoV-2

Hamsters were challenged at 4-5 weeks after the last dose with 1 × 10^4^ PFU of SARS-CoV-2 as described previously.^17^ In brief, hamsters in each group were divided into two cohorts and sacrificed three- or six-days post-challenge for viral load and pathology in lungs along with collection of nasal wash for upper respiratory viral load. Bodyweight and survival for each hamster was recorded daily post challenge until sacrifice. Euthanization, viral load, and histopathological examination were performed as described earlier.^17^

### Quantification of viral titer by cell culture infectious assay (TCID_50_)

The viral titer determination from lung tissue was performed as described previously. ^17^ In brief, the lungs were homogenized, clarified by centrifugation, and supernatant was diluted 10-fold and plated onto Vero cells in quadruplicate for live virus estimation. Similarly for nasal wash, the sample was centrifuged, diluted, and plated onto Vero cells. Cells were fixed, stained, and TCID_50_/mL was calculated by the Reed and Muench method.

### Real-time PCR for SARS-CoV-2 RNA Quantification

The SARS-CoV-2 RNA levels were measured using the established RT-PCR method to detect envelope gene of SARS-CoV-2 genome. RNA obtained from both lungs and nasal washes were analyzed for SARS-CoV-2 RNA levels as described previously. ^17, 19^

### Histopathology

As described previously, ^20, 21^ the left lungs of the hamsters were fixed with 4% paraformaldehyde for 1-week. The lungs were trimmed, processed, paraffin embedded, sectioned, and stained with Hematoxylin and Eosin (H&E) followed by microscopic scoring. The assessment of the pathological changes was done using scoring system that was used in the previous experiments where nine different areas of the lung sections are scored individually and averaged. In brief, a score of 0, was given to sections with no significant findings, score of 1 - for minor inflammation with slight thickening of alveolar septa and sparse monocyte infiltration, score of 2 - for apparent inflammation with alveolus septa thickening and interstitial mononuclear inflammatory infiltration, score of 3 and above - for diffuse alveolar damage with increased infiltration. ^17^

### Determination of serum and BAL S-2P specific IgG and IgA by ELISA

Serum and bronchoalveolar lavage samples (BAL) were evaluated for S-2P specific IgG and IgA antibody responses by ELISA. Briefly, 96-well Immulon 4HBX plates (Thermo Scientific, Cat# 3855) were coated with 1 μg/ml of S-2P, blocked using 5% BSA in PBS and, two-fold serially diluted serum or BAL samples were added onto the plate. Titers were determined using Sheep Anti-Mouse IgG-HRP (Jackson Immunoresearch, Cat # 515-035-071) or Rabbit Anti-Mouse IgA-HRP (Rockland, Cat # 610-4306). The endpoint titer (EPT) was determined by extrapolating from the closest OD values above and below the cutoff value (three times the mean background) and calculating the average of these two values.

### Neutralization Assays

The SARS-CoV-2 VSV pseudotype neutralization assay was performed at IBT Bioservices. In brief, the serum samples from mouse immunogenicity study were serially diluted two-fold, mixed with 10,000 RLU of rVSV-SARS-CoV-2 pseudovirus in which G gene of VSV is replaced with the firefly luciferase reporter gene and the S protein of SARS-CoV-2 is incorporated as the membrane protein on the surface of the VSV pseudotyped virus. The mixture was incubated at 37°C for 1 hour. Following incubation, the mixture was added to monolayer of Vero cells in triplicates and incubated for 24 hours at 37°C. After 24 h, firefly luciferase activity was detected using the Bright-Glo™ luciferase assay system (Promega Corporation, Cat # E2610). ID_50_ were calculated using XLfit dose response model.

The serum samples from hamsters were analyzed for neutralizing antibody titers using lentivirus expressing full-length wild type Wuhan-Hu-1 strain SARS-CoV-2 spike protein as described previously.^16^ Briefly, serum samples were heat-inactivated, serially diluted 2-fold in MEM with 2% FBS and mixed with equal volumes of pseudovirus. The samples were incubated at 37°C for 1 hour before adding to the HEK293-hACE2 plated cells. Cells were lysed 72 hours post incubation and relative luciferase units (RLU) were measured. ID50 and ID 90 (50% and 90% inhibition dilution titers) were calculated deeming uninfected cells as 100% and virus transduced control as 0%.

### Lung and Spleen Cytokine Assay

Lungs and spleens were dissected and manually disrupted to generate single-cell suspensions to be used in the Luminex and ELISpot assays. The contaminating red blood cells were lysed using 0.8% ammonium chloride with EDTA. The lymphocytes were washed with media, resuspended, and plated at 5 × 10^5^ cells per well in a 96-well flat bottom plate. The cells were stimulated with or without S-2P (5 μg/mL) and incubated at 37°C incubator with 5% CO_2_. After 72-hour incubation, the culture supernatants were collected and Luminex assay was performed according to the manufacturer’s protocol (EMD Millipore, Cat# MCYTOMAG-70K).

### Lung and Spleen B-cell ELISpot

Single cell suspensions from lungs and spleens of mice were stimulated with mouse IL-2 (R & D Systems, Cat # 402-ML; 0.5 μg/mL) and RD848 (Mabtech, Cat # 3611-5X; 1 μg/mL) for 3 days to induce nonspecific polyclonal expansion. At the end of 3 days, the cells were washed and plated onto PVDF ELISpot filter plates coated with anti-mouse IgG or IgA capture antibody (Mabtech, Cat# BASIC 3825-2H and BASIC 3835-2H). The plates were incubated at 37°C for 24 hours, following which the cells were stained with biotinylated S-2P antigen. Antigen-specific IgG- or IgA-producing B cells were detected using streptavidin-HRP. The spots were counted in AID ELISpot reader and expressed as spot forming units/million cells.

## Results

### Intranasal Immunization with S-2P/NE01 Induces Humoral Immune response in Mice

Data presented in **Figure 1** show a significant induction of serum S-2P-specific IgG after either intranasal or intramuscular vaccination. The route of vaccination did not impact seroconversion as all animals generated similar levels of anti-S-2P antibodies However, increased levels of IgA were detected only after intranasal vaccination, with no detectable levels of antigen-specific IgA in any of IM vaccinated animals. A cell-based neutralization assay utilizing an rVSV-pseudotype SARS-CoV-2 (**Table 1**), revealed that after 3 IN immunizations with S-2P/NE01, neutralizing antibodies were generated in the sera of all mice (8/8) with a GM IC_50_ >8000. Additionally, all mice (7/7) from the 2 IN S-2P/NE01 immunization group generated neutralizing antibodies but had a substantially lower GM IC_50_ of 1375.

**Figure 1.**
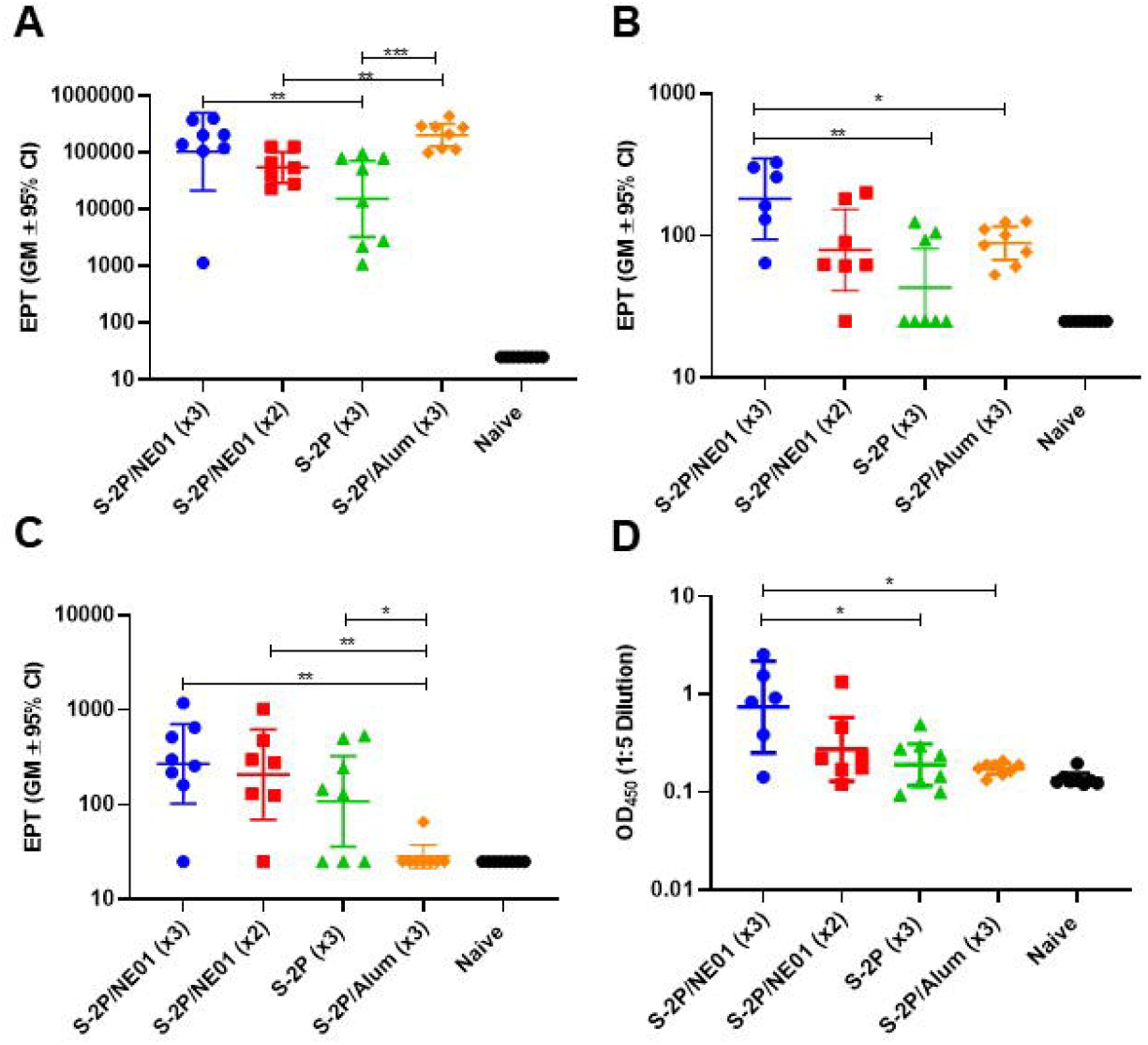
Humoral Immune Response in Mice Following Immunizations with S-2P. Humoral immune responses elicited in mice after immunizations with IN or IM formulations of S-2P as determined by A-B) Serum and BAL S-2P specific IgG End Point Titers (EPT). C) Serum IgA EPT and D) BAL IgA OD values. Statistical analysis was performed using Mann-Whitney nonparametric test for unpaired data, \**p<0.05*, \*\**p<0.01*, \*\*\**p<0.001*.

**Table 1.** Pseudovirus SARS-CoV-2 Neutralization activity of Serum from Mice

| **Groups** | **Vaccine** | **IC50 (Responders)** |
| --- | --- | --- |
| 1 | S-2P/NE01 (3X) | 8/8 (7/8 have IC50>8000) |
| 2 | S-2P/NE01 (2X) | 7/7 (1/7 have IC50>8000) |
| 3 | S-2P | 3/8 (0/8 have IC50>8000) |
| 4 | S-2P/Alum (3X) | 8/8 (6/8 have IC50>8000) |
| 5 | Naive | 0/8 |

Antibodies generated from 3 IM S-2P/Alum vaccinations had equivalent neutralizing activity to 3 IN immunizations.

### Intranasal Immunization Induces Mucosal Immunity in Mice

Mucosal immunity is defined by the induction of secretory IgA in mucosal surfaces and homing of immune cells to these tissues. Antigen-specific homing of B cells to mouse lungs and spleens were measured by ELISpot assay. There was a significant increase in homing of S-2P-specific IgG-producing B cells to the lungs after intranasal vaccination (2.5-fold increase in spot-forming units) and spleens (over 3.5-fold increase) compared to intramuscularly S-2P/alum vaccinated animals. In addition, only intranasal immunizations selectively produced B cells secreting S-2P-specific IgA in both spleens and lungs, suggesting a tissue resident memory B-cell response to the antigen, which supports a strong mucosal immune response conferred by this adjuvanted vaccine. (**Figure 2**).

**Figure 2.**
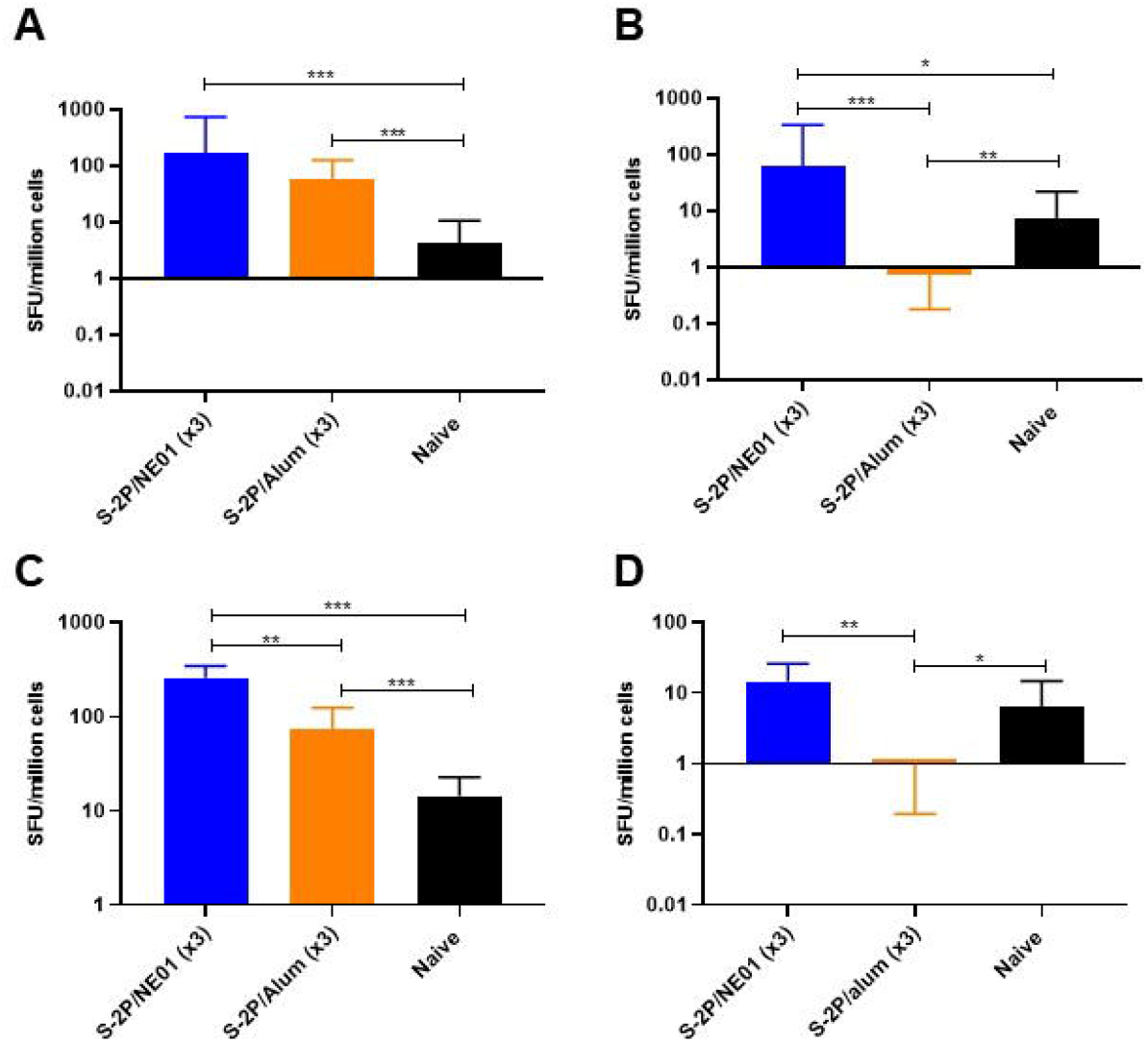
S-2P/NE01 Immunization Induces B- Cell Homing to Lungs and Spleen. Lungs and spleens were collected two weeks post last immunization and assessed for B cell homing by ELISpot. S-2P specific homing of IgG in lungs (A) or spleen (C): S-2P specific homing of IgA in lungs (B) or spleen (D). Data are presented as the geometric mean with a 95% confidence interval and statistical analysis was calculated using Mann-Whitney nonparametric test for unpaired data: \**p<0.05*, \*\**p<0.01*, \*\*\**p<0.001*.

### Balanced Th1/Th2 and Th17 Immune Response Induced by Intranasal Immunization in Mice

Cell-mediated immune responses were assessed in lung cells stimulated with S-2P antigen in a cytokine release assay. T_H_1 immune responses were evaluated by measuring IFNγ and TNFα production. IL-4 and IL-5 levels were used to assess T_H_2 responses. T_H_17 activity was measured by the release of IL-17A, the hallmark of mucosal immunity. As seen in **Figure 3A**, a significant induction of IFNγ was seen in lung tissue fromS-2P/NE01 IN immunized animals. These levels were statistically significant when compared to the levels in the lungs of S-2P/alum immunized mice. Similarly, an increased trend in TNF-α was also seen in IN immunized animals (data not shown). The T_H_2 immune response was significantly increased by both intranasal and intramuscular immunizations (**Figure 3B**). However, there was statistically significant induction of IL-4 in the lungs was seen in S-2P/alum (*p-value <0.05*) immunized mice compared to S-2P/NE01, although this result is not surprising as alum is known as a strong T_H_2 stimulating adjuvant. Mucosal immunity in the lungs was significantly stimulated by intranasal vaccination as evidenced by increased IL-17A levels in the immunized animals (**Figure 3C**): S-2P/NE01 immunized mice demonstrated more robust IL-17A with 100-fold higher increase compared to S-2P/alum immunized mice. Together, these data suggest that intranasal immunization with NE01-adjuvanted vaccine elicited a balanced Th1/Th2/Th17 immune response with production of tissue-resident memory T cells in the lung that will be beneficial for strong mucosal immunity.

**Figure 3.**
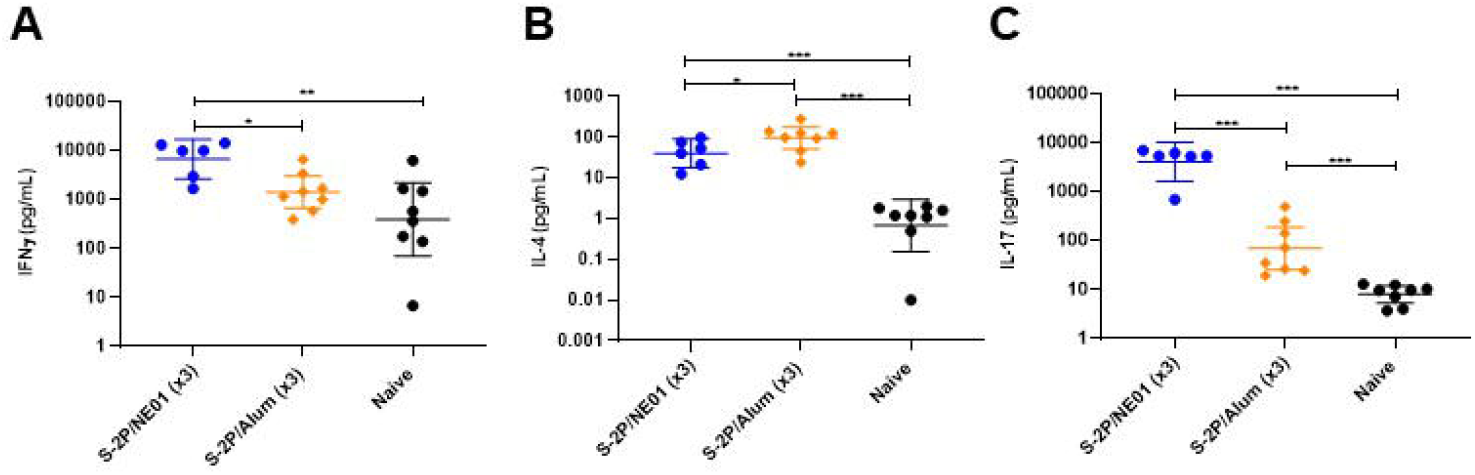
S-2P/NE01 Immunization Promotes Thl/Thl7 Cytokines in Lungs. Release ofINFγ(A), IL-4 (B): and IL-17(C) cytokines from S-2P stimulated single cell suspension of lungs. Data are presented as the geometric mean with a 95% confidence interval and statistical analysis was calculated between groups using Mann-Whitney nonparametric test for unpaired data, \**p<0.05*, \*\**p<0.01*, \*\*\**p<0.001*.

### Intranasal Immunizations Induce Highly Efficient Neutralizing Antibodies in Hamsters

To examine the vaccine efficacy of IN S-2P/NE01, a Syrian hamster model was selected due to SARS-CoV-2 pathogenesis and clinical symptoms of weight loss and fulminant pneumonia^22^. In this study, the same immunization protocol used in the mouse study was followed by dosing three weeks apart. Only the IN immunizations were performed in this study, comparing the efficacy of one versus two and three doses. Hamsters were challenged intranasally four weeks post last dose with 10^4^ PFU/hamster of SARS-CoV-2 isolate hCoV-19/Taiwan/4/2020. Animals were bled for serology prior to viral challenge to determine the systemic immune response. Although two different animal models were assessed, the convention of the immune response was similar as both models developed neutralizing antibodies after at least two doses. Statistically significant induction of neutralizing antibodies was seen in hamsters that received either two or three S-2P/NE01 immunizations with GMT for fifty-percent inhibition dose (ID_50_) at 825 after three IN vaccinations and 493 after two and GMT for ID90 at 195 and 104 respectively as assessed by pseudovirus neutralization assay. No induction of neutralizing antibodies was seen after one intranasal dose of the S-2P/NE01 vaccine (**Figure 4**).

**Figure 4.**
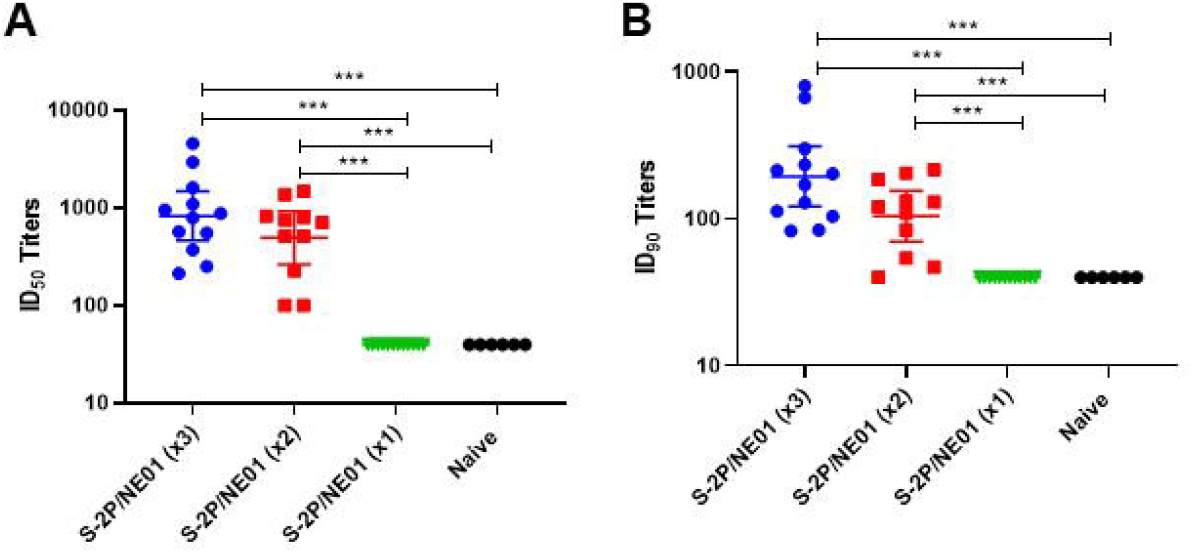
Neutralizing antibody titers from hamsters prior to challenge. Sera obtained from hamsters 10 days prior to challenge were analyzed by neutralization assay with pseudo\irus expressing SARS-CoV-2 spike protein to determine the ID_50_ (A) and ID_90_ (B) titers. Statistical significance between groups was calculated by one-way ANOVA: \**p<0.05*, \*\**p<0.01*, \*\*\**p<0.001*.

### Intranasal Immunizations Protect Hamsters from SARS-CoV-2 Challenge

Protection in the hamster challenge model is measured as a change in body weight after SARS-CoV-2 infection. In this study, hamsters that received either 2 or 3 IN doses of S-2P/NE01 gained between 1 and 2% of body weight measured every day until six days post-challenge. In contrast, animals immunized with 1 dose of S-2P/NE01 showed a weight loss similar to the control animals. Lung viral load at three- and six-days post-challenge measured by RT-PCR to detect viral RNA and by cell culture infectious assay (TCID50) showed a significant decrease in viral load in hamsters that received 2 or 3 IN doses of S-2P/NE01. Upper respiratory tract infection was measured in nasal washes collected at 3- and 6-days post-challenge. Both two- and three-dose immunized hamsters showed a two-fold decrease in viral load as measured by TCID_50_, 3-days post-challenge compared to control. However, six days post-challenge, the viral loads were below the limit of detection even in control. A significant decline in the number of copies of viral genome was observed six days post challenge in the nasal washes collected from three dose group (**Figure 5**). These results correlated with the bodyweight change and levels of neutralization antibodies, indicating two or three intranasal doses can provide protection to hamsters from both upper and lower respiratory tract infections bySARS-CoV-2.

**Figure 5.**
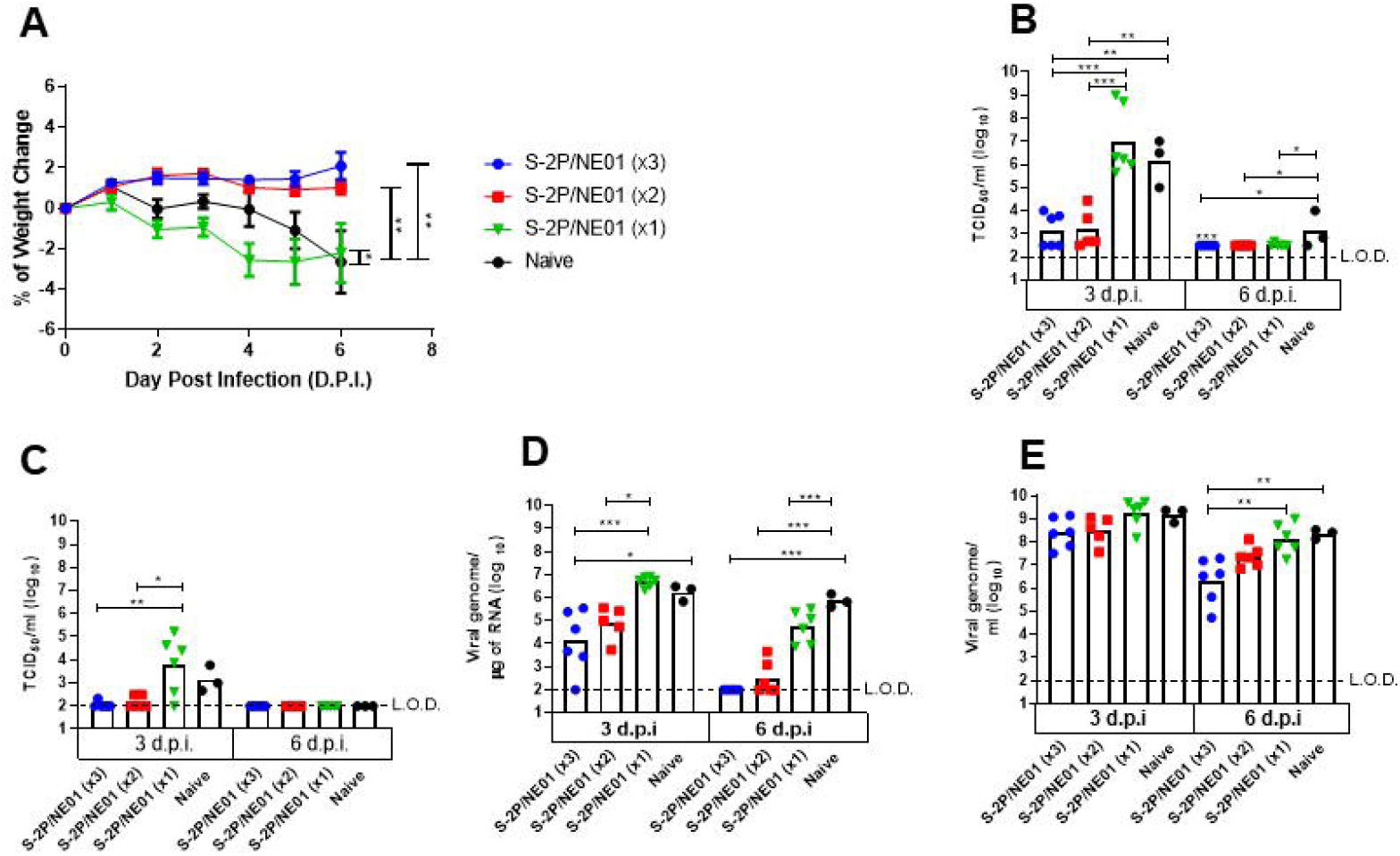
Intranasal S-2P/NE01 protects hamsters from SARS-CoV-2 Infection. Hamsters were challenged with10^4^ PFU of SARS-CoV-2 1-month post last immunization. Protection from infection is demonstrated by (A) daily measurement of body weights (B) viral load determination in lungs and (C) nasal wash by TCID50 and (D) quantitative PCR of λiral genome in lungs and (E) nasal wash. Dotted lines represent lower limit of detection. Statistical analysis for percent change in body weight was calculated with one-way ANOVA with Tukey’s multiple comparison test while \iral load by TCID50 and \iral genome was performed using Kruskal-Wallis with corrected Dunn’s multiple comparison test \**p*<0.05, \*\**p<0.01*, \*\*\**p<0.001*.

### Intranasal Immunizations Do Not Induce Lung Pathology

Lung sections from the hamsters were scored and analyzed for any pathological changes after infection. No differences in pathology were seen between the immunized groups and control after three days post-challenge. At 6 days post-challenge, animals immunized with either 2 or 3 times still had no detectable lung abnormalities, while animals in the control group and 1 dose immunized group showed significantly increased lung pathology with extensive immune cell infiltration and diffuse alveolar damage (**Figure 6 and Figure S1**). These results indicate that two or three doses of S-2P/NE01 vaccine induce a robust systemic immune response, in addition to local immunity, thereby enhancing viral clearance from lungs and nasal cavity and protecting hamsters from SARS-CoV-2 infection.

**Figure 6.**
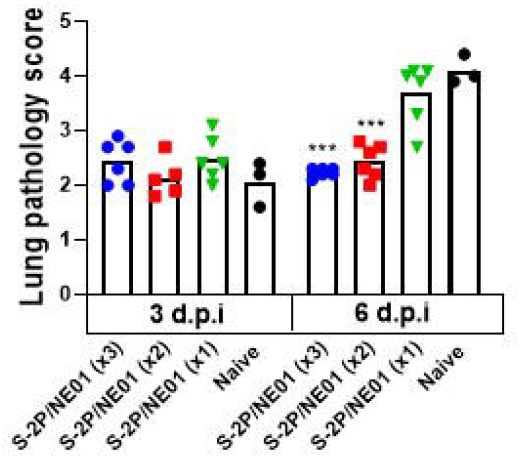
Intranasal S-2P/NE01 protects hamsters from lung pathology following infection with SARS-CoV-2. Hamsters were euthanized 3- and 6-dayspost challenge and lungs were collected for histopathological analysis. The lung sections were scored, and mean results are presented with error bars representing standard error. Statistical analysis calculated using one-way ANOVA with Tukey’s multiple comparison test.

## Discussion

Licensed SARS-CoV-2 vaccines had shown remarkable efficacy against infection and hospitalization. However, an increased rate of infections have been observed in vaccinated people contributing to the rise of a fourth and fifth wave of infections in countries that achieved high rates of vaccination post second or third immunizations. The rise of infections coincided with reduced SARS-CoV-2 antibody titers as well as spread of new variants of concern, especially the highly contagious Omicron (BA.1) variant in addition to other localized variants: Alpha (B.1.1.7), Beta (B.1.351), Gamma (P1), and Delta (B.1.617.1). Immune evasion can be observed with these variants by antigenic drift in the receptor-binding domain leading to reduced efficacy of vaccine-induced neutralizing antibodies. In spite of these observations, all COVID-19 vaccines still exhibit high efficacy against hospitalization and severe disease. ^23, 24^ Administration of a booster dose to those vaccinated six months or more following the last dose of vaccination is proposed as a remedy to boost serum antibodies which in turn could reduce SARS-CoV-2 infections and transmission, ^25^ thus reducing chances for the emergence of new variants. We believe that any proposed solution based on boosting serum antibodies by administration of a third vaccination to influence nasal colonization and spread of the virus is a temporary solution as intramuscular immunization does not elicit mucosal immunity, the only permanent and efficient solution to the problem.

Our mucosal adjuvant NE01 demonstrates a potential long-lasting induction of mucosal and systemic immunity, achieved by an intranasal administration of a NE01 formulated/adjuvanted vaccine. Intranasal vaccination using nasal NE01 adjuvant/delivery had shown unique attributes including elicitation of mucosal Th17, IgA, serum IgG, and homing of IgG and IgA B- and T-cells to reside in mucosal tissues. These attributes were absent when vaccines delivered intramuscularly. In addition, our adjuvant induces IL-17. Current clinical evidence has shown that Th17 polarization in COVID-19 patients can be associated with poor disease outcomes facilitated by eosinophilic infiltrates in the lungs.^26^ However, NE01-intranasal vaccines has been previously evaluated in primary animal models for RSV (cotton rats) and pandemic flu (ferrets) eliciting mucosal and systemic immunity that not only prevented disease, but also prevented nasal colonization following intranasal and intratracheal viral challenge. ^27, 28^ In these studies, local, but not systemic increases of IL-17 were observed in the lung without co-expression of IL-13, where IL-13 has been associated with severe disease progression with COVID-19 in mouse models.^29^ Moreover, pre-clinical mouse studies using nanoemulsion-inactivated RSV demonstrated no immunopotentiation, with absence of mucus hypersecretion and lack of airway eosinophilia. ^30^ Since our vaccine platform consistently contributes to balanced T-cell immunity (Th1/Th2/Th17), skewed and potentially damaging T-cell polarizations are likely negated due to NE01’s unique adjuvant mechanism of action that induces homing of memory cells and induction of mucosal immunity at distant mucosal tissues. Our previously reported data showed that intranasal immunization with a bivalent gD2/gB2/NE01 vaccine elicited mucosal immunity that prevented colonization and infection following intravaginal HSV2 challenge in a guinea pig model. ^31^ Data presented in the current study show that formulation of SARS-CoV-2 S-2P antigen in NE01 elicited protective immune responses against lung infection and disease evidenced by histopathologic scoring. Further, intranasally vaccinated animals exhibited an enhanced reduction of SARS-CoV-2 viral load in the lungs and nasal washes. With the caveat that IM vaccination temporarily reduced nasal colonization following vaccination, our intranasal vaccination outcomes were in line with other data generated in the same hamster model using an S-2P vaccine adjuvanted with a combination of Alum and CpG 1018,^16, 17^ suggesting that intranasal immunization could be as efficient as intramuscular vaccination with the potential advantage of induction of mucosal immunity that would eliminate the virus at its port of entry.

The NE01 adjuvant is a clinical-stage adjuvant and has been evaluated in several clinical trials, including a phase 1 anthrax vaccine trial and a seasonal flu trial.^32^ NE01-adjuvanted vaccines demonstrated a remarkable safety profile and a robust mucosal and systemic immunity. Additionally, the exceptional stability (at 5°C) and ease of administration, reduces the complexities involved with ultra-low cold chain storage and needle-less administration, making this vaccine attractive to low-income countries. We believe our NE01 technology can play a role in providing safe and efficacious standalone vaccine to protect against infection and disease. In the light of the fact that billions of people had already received IM vaccines and that many vaccines are already licensed and have been used, our future development plan includes using this unique intranasal vaccine as a booster vaccine to those who had received IM vaccines fin order to boost their systemic immunity and to confer complementary mucosal immunity to achieve the ultimate goal of eliciting immunity for the prevention of colonization, spread, infection, and disease caused by SARS-CoV2.

## Acknowledgements

We are grateful for the participation of Dr. Han van den Bosch for manuscript review and constructive comments. We also thank team members at TFBS Bioscience Incorporation for hamster housing and immunization process. We thank the Biomedical Translation Research Center, Academia Sinica, Taiwan, for performing hamster challenge. In addition, we would like to acknowledge Dr. Yu-Chi Chou and his team at the RNAi Core Facility, Academia Sinica for the pseudovirus neutralization assay.

## Data availability

The datasets generated in this manuscript are available from the corresponding author on reasonable request.

## References

1. Coronaviridae Study Group of the International Committee on Taxonomy of V. The species Severe acute respiratory syndrome-related coronavirus: classifying 2019-nCoV and naming it SARS-CoV-2. Nat Microbiol 2020; 5:536–44.

2. Liu Y, Ning Z, Chen Y, Guo M, Liu Y, Gali NK, et al. Aerodynamic analysis of SARS-CoV-2 in two Wuhan hospitals. Nature 2020; 582:557–60.

3. Sungnak W, Huang N, Becavin C, Berg M, Queen R, Litvinukova M, et al. SARS-CoV-2 entry factors are highly expressed in nasal epithelial cells together with innate immune genes. Nat Med 2020; 26:681–7.

4. Lee IT, Nakayama T, Wu CT, Goltsev Y, Jiang S, Gall PA, et al. ACE2 localizes to the respiratory cilia and is not increased by ACE inhibitors or ARBs. Nat Commun 2020; 11:5453.

5. Hou YJ, Okuda K, Edwards CE, Martinez DR, Asakura T, Dinnon KH, 3rd, et al. SARS-CoV-2 Reverse Genetics Reveals a Variable Infection Gradient in the Respiratory Tract. Cell 2020; 182:429–46 e14.

6. Gallo O, Locatello LG, Mazzoni A, Novelli L, Annunziato F. The central role of the nasal microenvironment in the transmission, modulation, and clinical progression of SARS-CoV-2 infection. Mucosal Immunol 2021; 14:305–16.

7. Subbarao K, Mahanty S. Respiratory Virus Infections: Understanding COVID-19. Immunity 2020; 52:905–9.

8. Prevention CfDCa. Science Brief: COVID-19 Vaccines and Vaccination. https://www.cdc.gov/coronavirus/2019-ncov/science/science-briefs/fully-vaccinated-people.html.

9. Prevention CfDCa. CDC Real-World Study Confirms Protective Benefits of mRNA COVID-19 Vaccines. https://www.cdc.gov/media/releases/2021/p0329-COVID-19-Vaccines.html, 2021.

10. Polack FP, Thomas SJ, Kitchin N, Absalon J, Gurtman A, Lockhart S, et al. Safety and Efficacy of the BNT162b2 mRNA Covid-19 Vaccine. N Engl J Med 2020; 383:2603–15.

11. Baden LR, El Sahly HM, Essink B, Kotloff K, Frey S, Novak R, et al. Efficacy and Safety of the mRNA-1273 SARS-CoV-2 Vaccine. N Engl J Med 2021; 384:403–16.

12. Sadoff J, Gray G, Vandebosch A, Cardenas V, Shukarev G, Grinsztejn B, et al. Safety and Efficacy of Single-Dose Ad26.COV2.S Vaccine against Covid-19. N Engl J Med 2021; 384:2187–201.

13. Knoll MD, Wonodi C. Oxford–AstraZeneca COVID-19 vaccine efficacy. The Lancet 2021; 397:72–4.

14. Pearson CF, Jeffery R, Oxford-Cardiff C-LC, Thornton EE. Mucosal immune responses in COVID19 - a living review. Oxf Open Immunol 2021; 2:iqab002.

15. Russell MW, Moldoveanu Z, Ogra PL, Mestecky J. Mucosal Immunity in COVID-19: A Neglected but Critical Aspect of SARS-CoV-2 Infection. Front Immunol 2020; 11:611337.

16. Kuo TY, Lin MY, Coffman RL, Campbell JD, Traquina P, Lin YJ, et al. Development of CpG-adjuvanted stable prefusion SARS-CoV-2 spike antigen as a subunit vaccine against COVID-19. Sci Rep 2020; 10:20085.

17. Lien CE, Lin YJ, Chen C, Lian WC, Kuo TY, Campbell JD, et al. CpG-adjuvanted stable prefusion SARS-CoV-2 spike protein protected hamsters from SARS-CoV-2 challenge. Sci Rep 2021; 11:8761.

18. Makidon PE, Bielinska AU, Nigavekar SS, Janczak KW, Knowlton J, Scott AJ, et al. Pre-clinical evaluation of a novel nanoemulsion-based hepatitis B mucosal vaccine. PLoS One 2008; 3:e2954.

19. Corman VM, Landt O, Kaiser M, Molenkamp R, Meijer A, Chu DK, et al. Detection of 2019 novel coronavirus (2019-nCoV) by real-time RT-PCR. Euro Surveill 2020; 25.

20. Liu L, Wei Q, Lin Q, Fang J, Wang H, Kwok H, et al. Anti-spike IgG causes severe acute lung injury by skewing macrophage responses during acute SARS-CoV infection. JCI Insight 2019; 4.

21. Jiang RD, Liu MQ, Chen Y, Shan C, Zhou YW, Shen XR, et al. Pathogenesis of SARS-CoV-2 in Transgenic Mice Expressing Human Angiotensin-Converting Enzyme 2. Cell 2020; 182:50–8 e8.

22. Le Bras A. Development of a new preclinical model of Parkinson’s disease. Lab Animal 2020; 49:219-.

23. Lopez Bernal J, Andrews N, Gower C, Gallagher E, Simmons R, Thelwall S, et al. Effectiveness of Covid-19 Vaccines against the B.1.617.2 (Delta) Variant. N Engl J Med 2021.

24. Lustig Y, Zuckerman N, Nemet I, Atari N, Kliker L, Regev-Yochay G, et al. Neutralising capacity against Delta (B.1.617.2) and other variants of concern following Comirnaty (BNT162b2, BioNTech/Pfizer) vaccination in health care workers, Israel. Euro Surveill 2021; 26.

25. Administration FaD. FDA Authorizes Booster Dose of Pfizer-BioNTech COVID-19 Vaccine for Certain Populations. https://www.fda.gov/news-events/press-announcements/fda-authorizes-booster-dose-pfizer-biontech-covid-19-vaccine-certain-populations, 2021.

26. Hotez PJ, Bottazzi ME, Corry DB. The potential role of Th17 immune responses in coronavirus immunopathology and vaccine-induced immune enhancement. Microbes Infect 2020; 22:165–7.

27. O’Konek JJ, Makidon PE, Landers JJ, Cao Z, Malinczak CA, Pannu J, et al. Intranasal nanoemulsion-based inactivated respiratory syncytial virus vaccines protect against viral challenge in cotton rats. Hum Vaccin Immunother 2015; 11:2904–12.

28. Hamouda T, Sutcliffe JA, Ciotti S, Baker JR, Jr. Intranasal immunization of ferrets with commercial trivalent influenza vaccines formulated in a nanoemulsion-based adjuvant. Clin Vaccine Immunol 2011; 18:1167–75.

29. Donlan AN, Sutherland TE, Marie C, Preissner S, Bradley BT, Carpenter RM, et al. IL-13 is a driver of COVID-19 severity. JCI Insight 2021; 6.

30. Lindell DM, Morris SB, White MP, Kallal LE, Lundy PK, Hamouda T et al. A novel inactivated intranasal respiratory syncytical virus vaccine promotes viral clearance without Th2 associated vaccine-enhanced disease. PLoS One 2011; 6(7)

31. Bernstein DI, Cardin RD, Bravo FJ, Hamouda T, Pullum DA, Cohen G, et al. Intranasal nanoemulsion-adjuvanted HSV-2 subunit vaccine is effective as a prophylactic and therapeutic vaccine using the guinea pig model of genital herpes. Vaccine 2019; 37:6470–7.

32. Stanberry LR, Simon JK, Johnson C, Robinson PL, Morry J, Flack MR, et al. Safety and immunogenicity of a novel nanoemulsion mucosal adjuvant W805EC combined with approved seasonal influenza antigens. Vaccine 2012; 30:307–16.

