## Supplementary figures and images for "Intranasal Nanoemulsion Adjuvanted S-2P Vaccine Demonstrates Protection in Hamsters and Induces Systemic, Cell-Mediated and Mucosal Immunity in Mice"

### Supplemental Figure S1

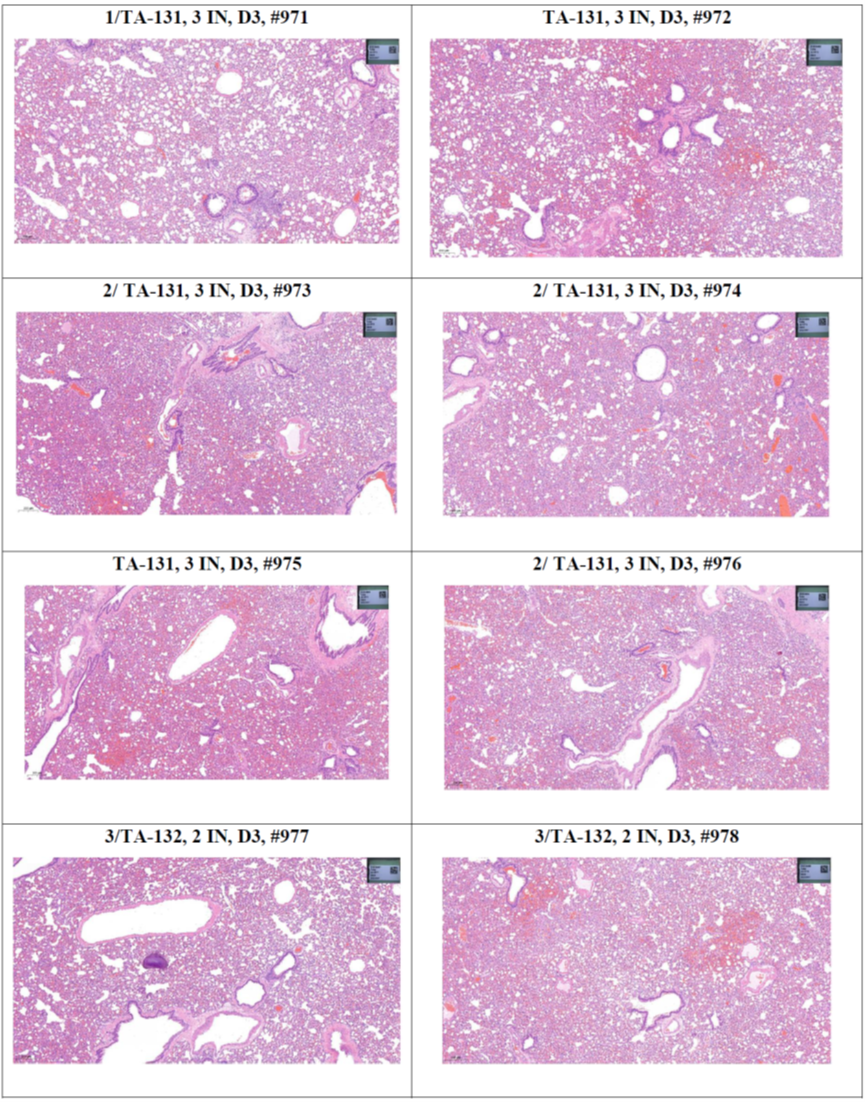

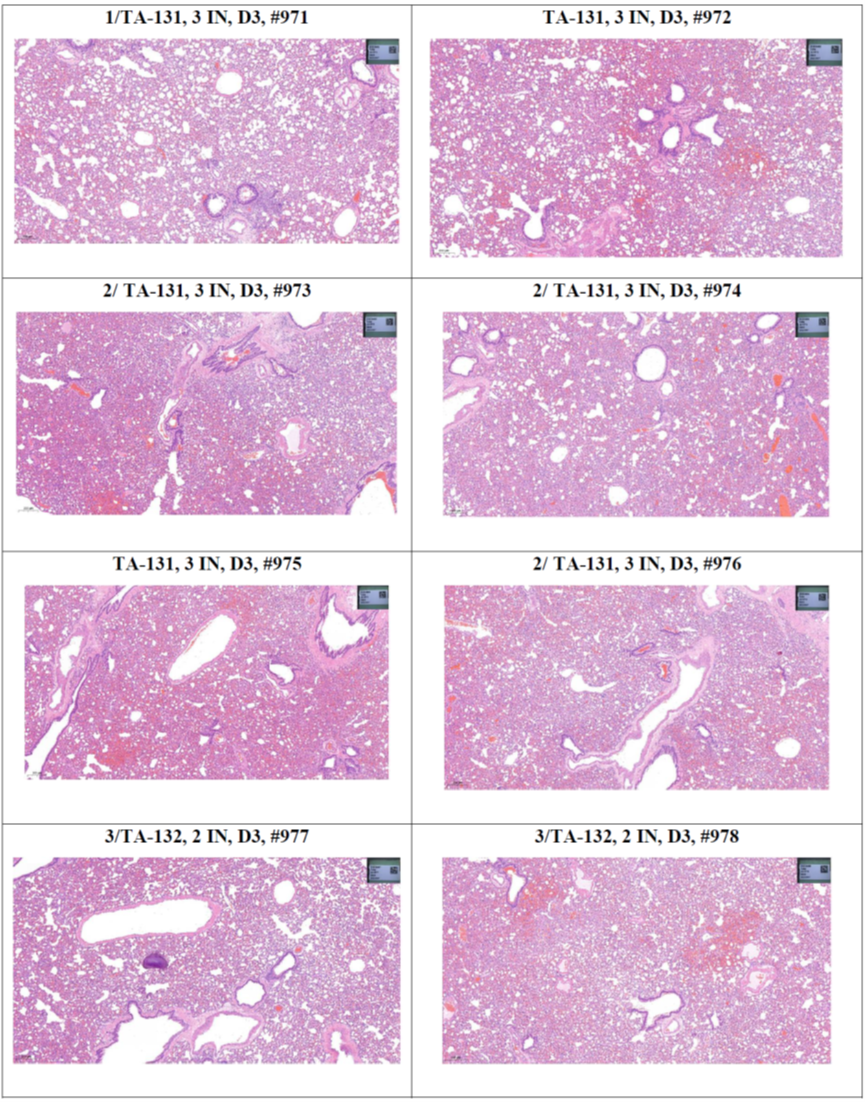

**3 dpi	6 dpi**

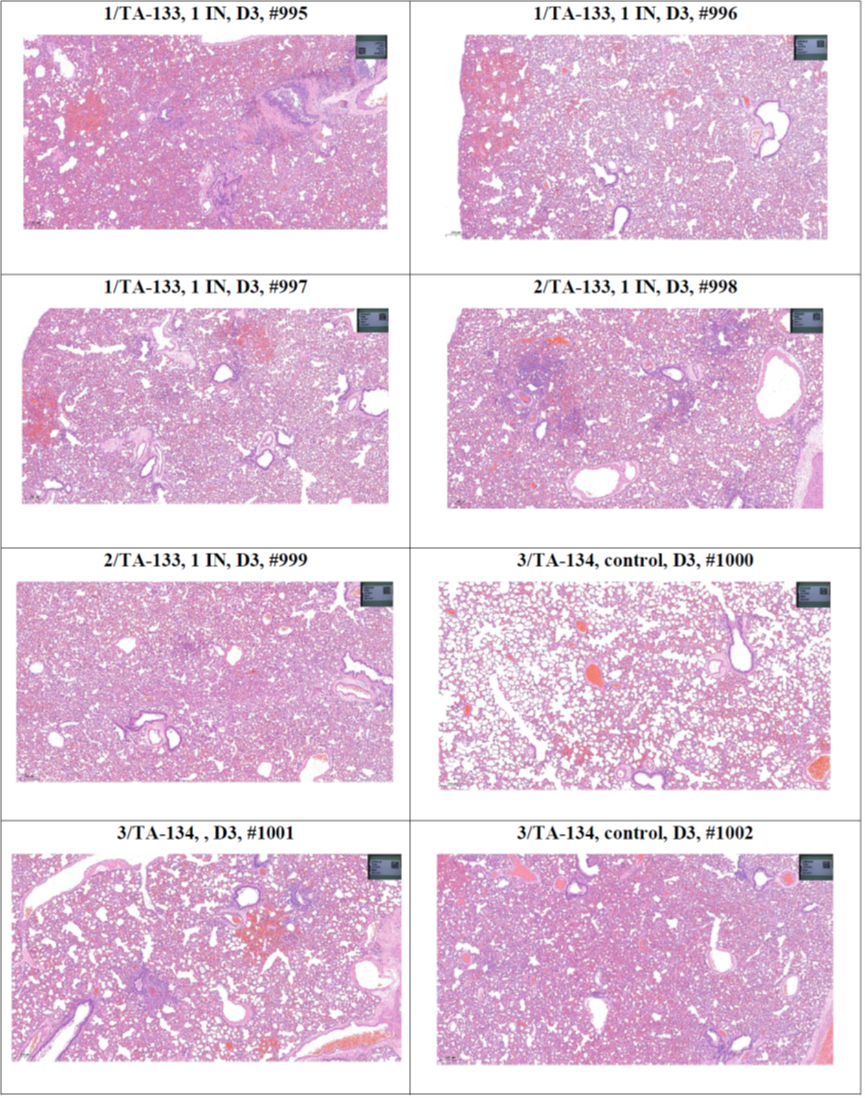

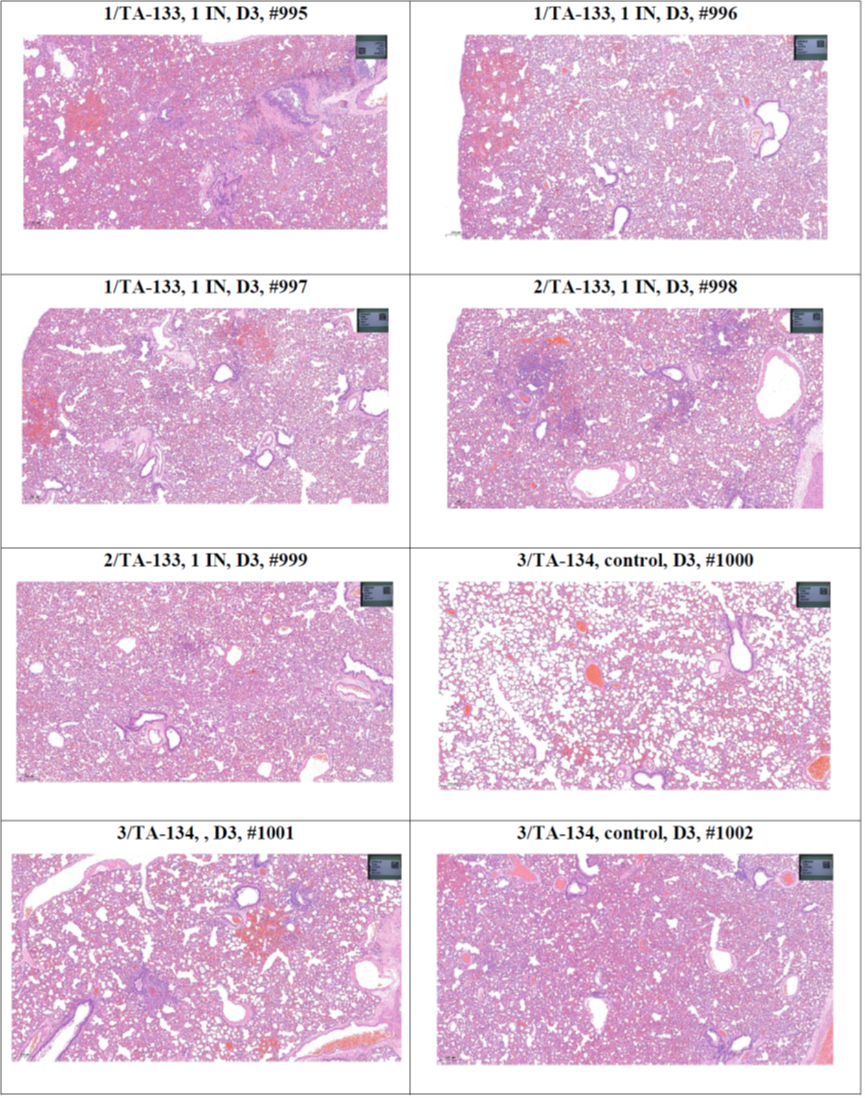

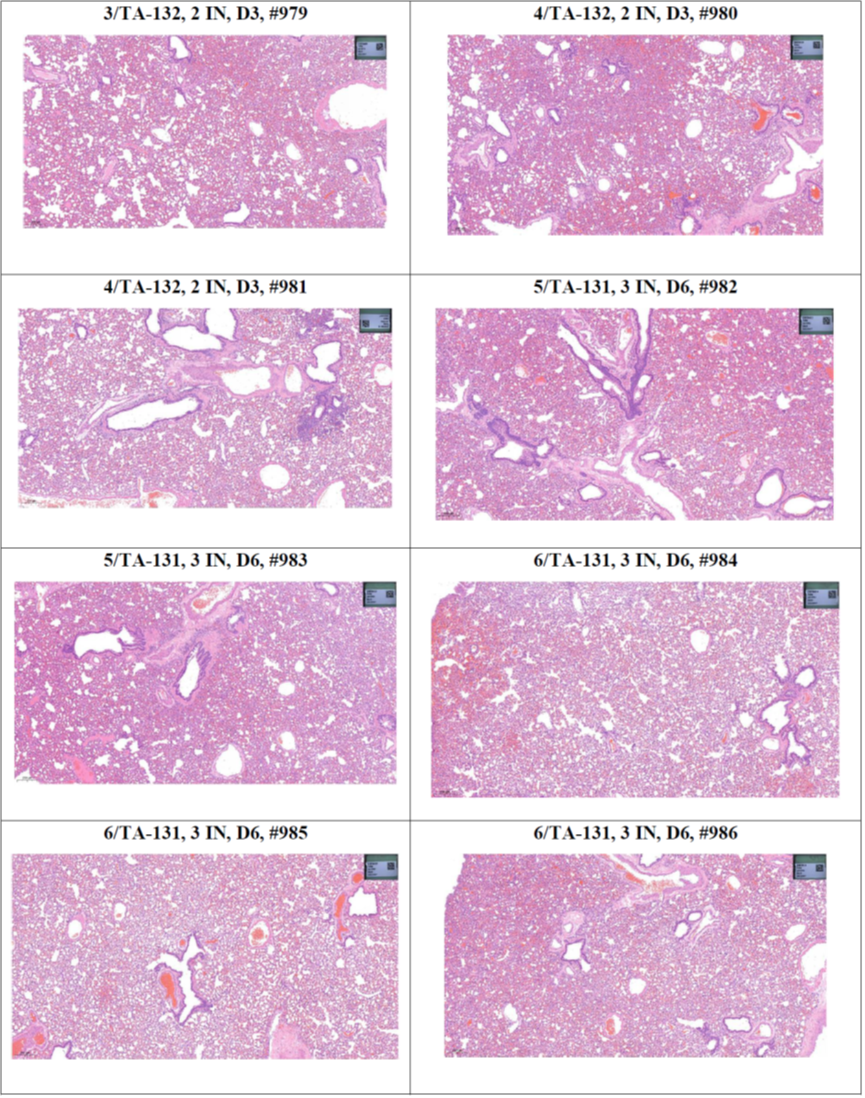

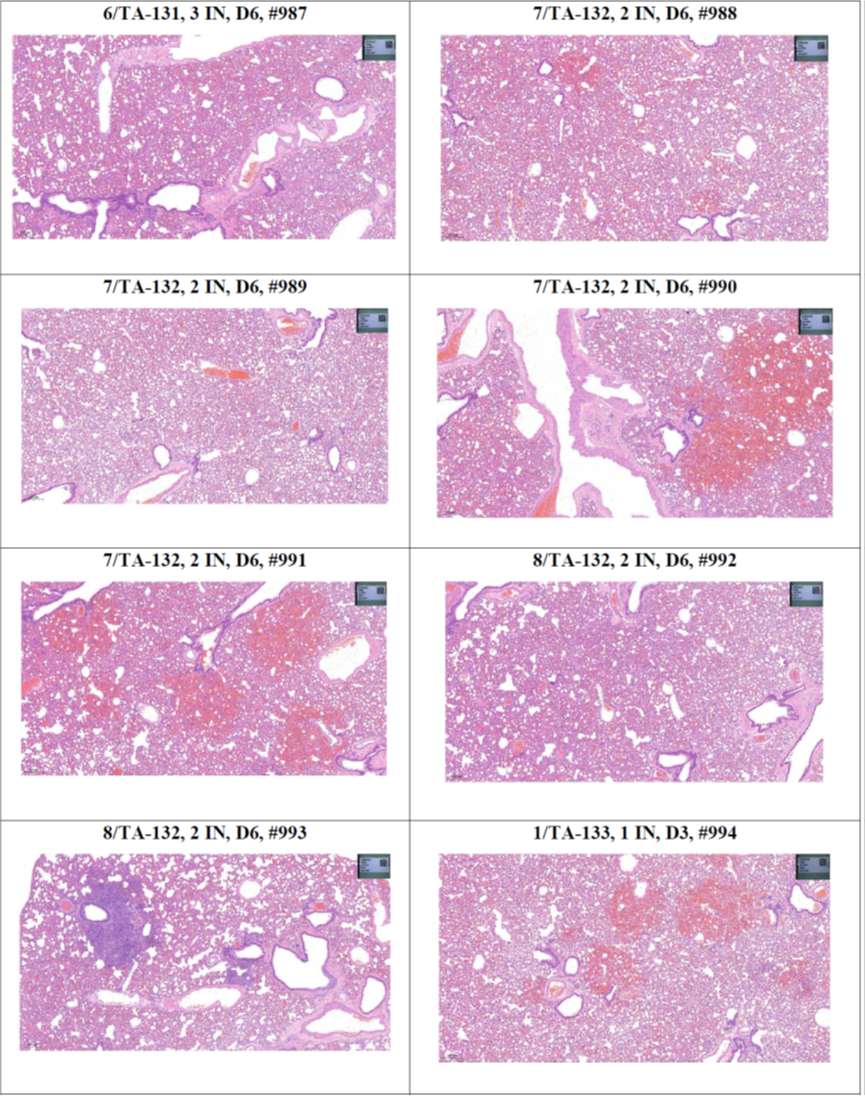

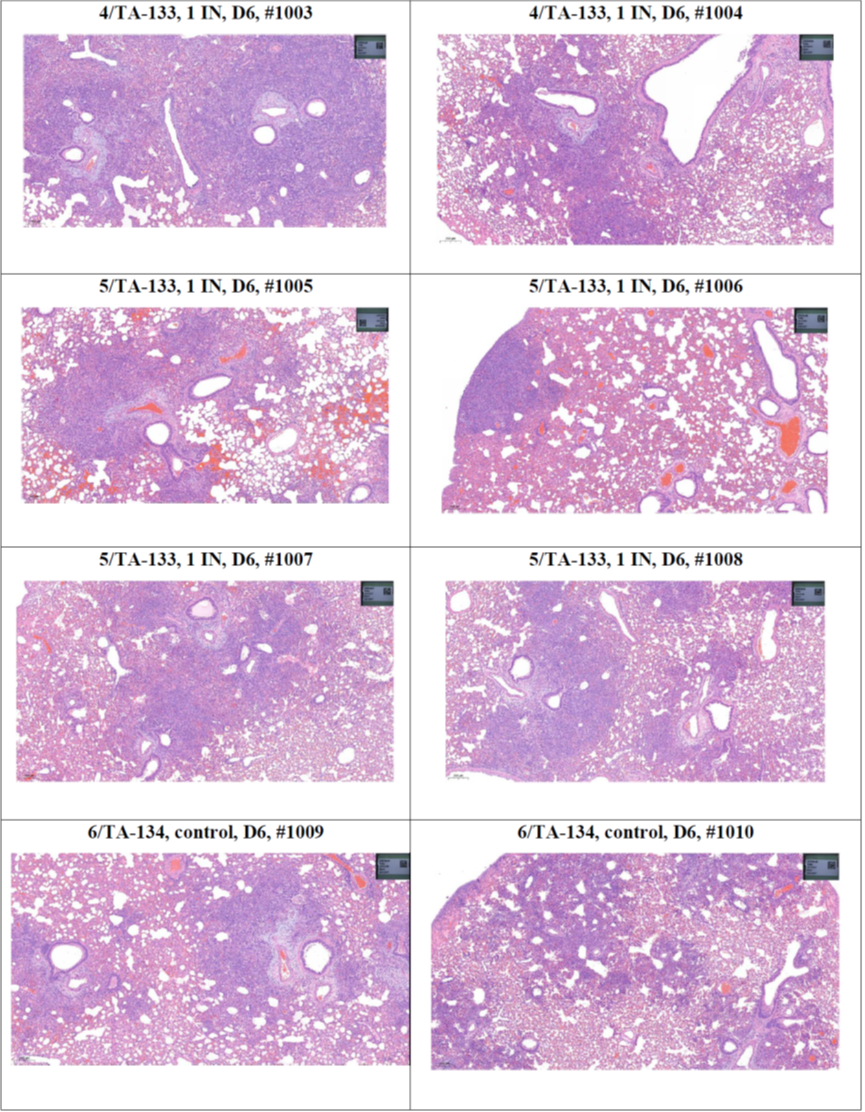

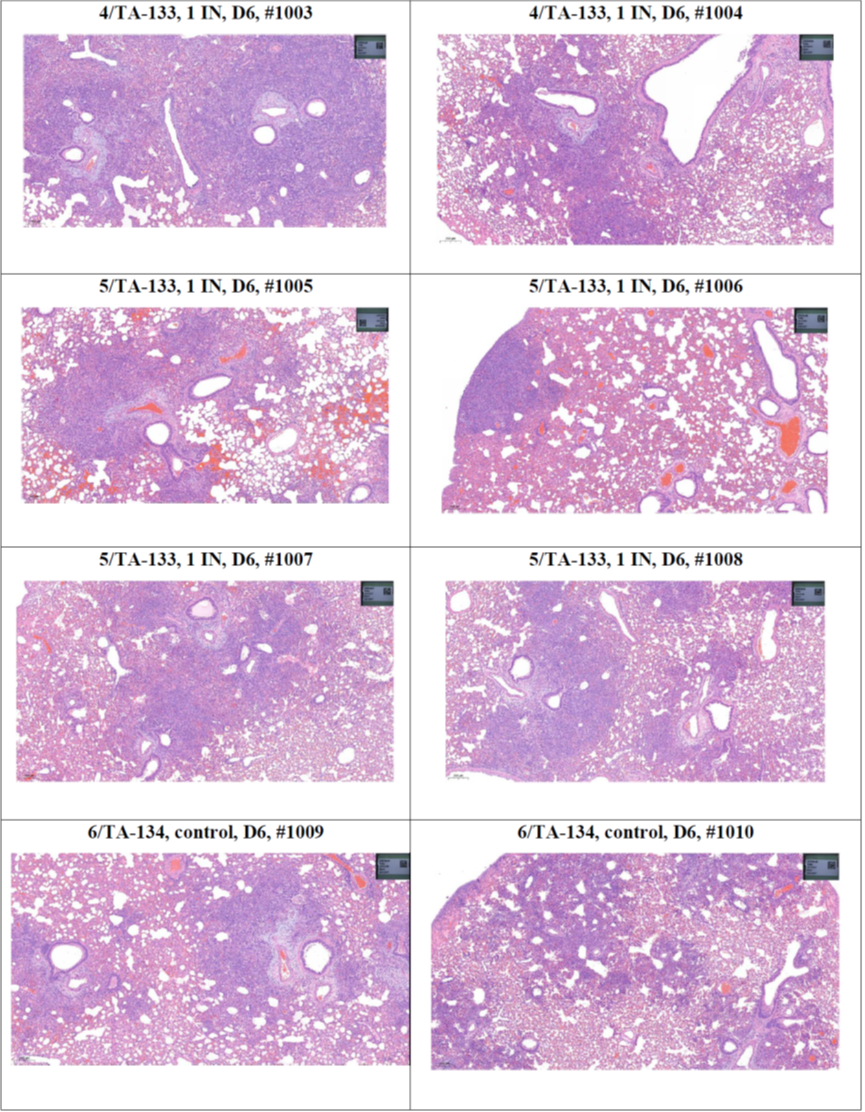

**S-2P/NE01 (x3)**

**S-2P/NE01 (x2)**

**S-2P/NE01 (x1)**

**Naive**
